# Sex-specific metabolic responses to modulation of the glucagon-FGF21 axis in female mice

**DOI:** 10.1101/2025.04.09.647697

**Authors:** Christoffer Merrild, Valdemar Brimnes Ingemann Johansen, Christoffer Clemmensen, Pablo Ranea-Robles

## Abstract

Fibroblast growth factor 21 (FGF21) and glucagon are key regulators of energy homeostasis. However, male-biased preclinical studies overlook critical sex differences in metabolism, hindering our understanding of FGF21’s role in glucagon’s effects on females and effective treatments for women with metabolic diseases. We investigated the physiological effects of FGF21 deficiency in female mice fed a chow or obesogenic diet. During chow-feeding, FGF21 deficiency had minimal impact on body weight, response to fasting, and voluntary exercise. However, Fgf21 knockouts (KO) fed the obesogenic diet exhibited increased adiposity compared to wild-type females. Long-acting glucagon analog treatment (LA-Gcg) reduced body weight in both genotypes but exacerbated glucose intolerance in female KOs. We also compared the effects of LA-Gcg and semaglutide, a glucagon-like peptide 1 analog, in diet-induced obese males and females. LA-Gcg treatment induced greater weight loss and reduced food intake in males, whereas semaglutide had similar effects across sexes. Furthermore, glucose tolerance was significantly worse in LA-Gcg-treated females compared to males. Concluding, FGF21 deficiency potentiates diet-induced obesity in female mice, and LA-Gcg elicits sex-specific effects on body weight and glucose tolerance, highlighting potential sexual dimorphisms to glucagon-based therapies and underscoring the importance of considering sex as a biological variable in metabolic research.

**Highlights:**

- Biological sex fundamentally affects metabolism, yet this variable remains largely underexplored in metabolic research.
- This study investigates how FGF21 deficiency affects metabolic responses in female mice, and compares the effects of a glucagon analog (LA-Gcg) and semaglutide in both sexes.
- FGF21 deficiency exacerbates diet-induced obesity in female mice. LA-Gcg’s effects on body weight and glucose tolerance are modulated by FGF21 and sex-dependent, with females showing a blunted response to weight loss and worse glucose tolerance.
- Our findings highlight sex-specific differences in metabolic responses, emphasizing the need to consider sex as a key variable in the development of glucagon-based therapies.

## Introduction

Human metabolism is deeply influenced by biological sex^1^, with numerous physiological processes exhibiting sexual dimorphism. This influence extends to preclinical models, where murine studies have revealed sex-specific variations in various metabolic parameters^2^. Nevertheless, a persistent bias toward male rodents in metabolic research has created a significant knowledge gap regarding female-specific responses^3–6^, particularly in the context of obesity and its associated comorbidities^7^. This oversight is especially worrisome given the near-equal prevalence of obesity in both sexes^8,9^ and the documented sex-dependent differences in its related complications^10^, including a male predominance in diabetes and a female predominance in severe obesity^11–13^.

Fibroblast growth factor 21 (FGF21) and glucagon, both essential regulators of energy homeostasis, have been extensively studied and proposed as potential therapeutic targets for obesity and diabetes^14,15^. FGF21 is a hormone with multifaceted roles in energy, lipid, and glucose homeostasis^14,16,17^. It is primarily secreted by hepatocytes in response to various stressors, such as acute fasting, protein restriction, and a ketogenic diet in rodent models^16,18,19^. In the liver, FGF21 primarily reduces *de novo* lipogenesis and enhances fatty acid beta-oxidation, thereby protecting against lipotoxicity^14^. Research on FGF21’s actions has primarily focused on male mice^14^, leaving its role in female physiology, apart from reproductive function^20^, largely unexplored. However, several recent studies have uncovered sex-specific differences in FGF21’s metabolic regulation, including responses to protein restriction^21,22^, energy homeostasis^23,24^, calorie-restriction-induced beiging of white adipose tissue^25^, and hepatic lipid metabolism^26,27^. These findings highlight the relevant sexual dimorphism in FGF21-mediated metabolic functions and stress the necessity for more comprehensive sex-specific studies to fully understand FGF21 biology.

The hormone glucagon plays a key role in glucose and energy homeostasis^15,28^. While pharmacological antagonism of the glucagon receptor has been explored for glycemic management^29,30^, it is associated with adverse cardiovascular events^15^. Conversely, glucagon receptor agonism is now being investigated as a therapeutic strategy for obesity management in combination with incretin analogs^31^, potentially converging on FGF21 signaling^32^. Indeed, preclinical studies in male mice suggest that FGF21 is vital for glucagon’s beneficial metabolic effects on weight loss and lipid metabolism^32,33^, with evidence showing that glucagon increases FGF21 secretion^34^. Despite reported sex differences in pancreatic islet biology, liver metabolism^35,36^, and in the efficacy of the triple GIP-GLP1-GCG agonist Retatrutide (LY3437943; Eli Lilly)^31^, very little is known about sex-specific differences in glucagon biology. In addition, whether FGF21 mediates glucagon’s beneficial effects in female mice has not been investigated.

Given the therapeutic potential of FGF21 and glucagon, and the observed sexual dimorphism in these metabolic pathways, understanding the glucagon-FGF21 axis in females is important. Therefore, in this study, we investigated the physiological effects of FGF21 deficiency in female mice fed a chow diet and on an obesogenic diet. Furthermore, we explored potential sex-specific differences in response to glucagon agonism by comparing its effects in male and female mice.

## Research Design and Methods

### Animals and housing conditions

Fibroblast growth factor 21 (Fgf21) knockout (KO) mice (Fgf21*^tm^*^1^*^.1Djm^*, The Jackson Laboratory, stock number 033846 available at the CBMR, RRID:IMSR_JAX:033846) on a C57BL/6J background (not verified) were used. Wild-type (WT) and Fgf21 KO littermates were generated by breeding Fgf21 heterozygous mice. Offspring was genotyped using protocol 44140 recommended by Jackson Labs. For the semaglutide and glucagon head-to-head study, male and female C57BL/6J mice were purchased from Janvier and fed a high-fat, high-sucrose diet as explained below (HFHS, Research Diets, #D12331i). Mice were kept in individually ventilated cages at an ambient temperature of 22°C ± 2°C, with humidity ranging from 45% to 65%. They were exposed to 12 hours of light (06:00-18:00) followed by a 12-hour dark phase (18:00-06:00) and had *ad libitum* access to water and either a chow diet (Safe Diets, #Safe D30) or a HFHS diet (Research Diets, D12331i, 5.56 kcal/g: 58% fat, 25.5% carbohydrate, and 16.4% protein). All female mice were housed in pairs during the experimental procedures, while male mice were housed individually. All mouse experiments were performed according to the ARRIVE guidelines^37^ and were approved by the Danish Animal Experimentation Inspectorate (2018-15-0201-01457 and 2023-15-0201-01442).

### Metabolic phenotyping of WT and Fgf21 KO female mice

The first cohort of female mice (cohort 1; n=20 WT, n=19 Fgf21 KO; age=12.5 months) was phenotyped while they had *ad libitum* access to a chow diet (Safe Diets, #SAFE D30) for 37 days. Body weight and food intake were monitored throughout the study period, while body mass composition was assessed after 37 days (for details, see below). Subsequently, mice were subjected to overnight (16 hours) fasting. Food was removed from the cage grids at 18:00, cage bedding was replaced, and mice were food-deprived until the next day at 10:00 am. Before food was placed back into the cage grid, mice were weighed, blood beta-hydroxybutyrate (β-OHB) was measured (Blood β-Ketone Test Strips, Precision, Abbott) from a drop of tail blood, and a blood sample was collected from the tail vein to obtain plasma for further biochemical measurements. This initial cohort of mice was then switched to *ad libitum* access to a HFHS diet (Research Diets, #D12331i). After 94 days of HFHS feeding, mice were subjected to an intraperitoneal glucose tolerance test (ipGTT), and on day 112 their body composition was measured as explained below. A total of n=19 WT and n=17 Fgf21 KO mice entered the HFD-feeding, where 6 WT and 7 Fgf21 KO mice were excluded/euthanized due to overgrooming, tumors, or sudden death. A second cohort of 12-month-old female mice (cohort 2; n=7 WT, n=6 Fgf21 KO) was exposed to voluntary running for 31 days. Running distance was monitored with a cyclometer and annotated three times a week. Two WT mice in cohort 2 were excluded from the study since they did not engage in voluntary running. After the examination of voluntary running, mice were euthanized, and blood and tissues were collected.

### *In vivo* glucagon pharmacology in diet-induced obese WT and Fgf21 KO female mice

A long-acting glucagon analog (LA-Gcg, Novo Nordisk Compound Sharing, #NNC9204-0043) was dissolved in a vehicle solution consisting of 145 mM propylene glycol (Sigma-Aldrich, #398039) and 50 mM sodium phosphate buffer at pH=7.4. Vehicle or LA-Gcg was administered daily subcutaneously to diet-induced obese WT and Fgf21 KO female mice from cohort 1 (for details, see above), close to the dark phase, at 5 µL/g body weight to obtain a 30 nmol/kg dose for 16 days. On experimental day 13, mice were subjected to an ipGTT, as detailed below. On day 14, their body compositions were analyzed using MRI (for details, see below). Body weight and food intake were measured daily at the time of daily injection, just before the dark phase. Mice were euthanized on day 16 and blood and tissues were collected. Exclusions: Two WT vehicle-treated mice (one died suddenly, and one was excluded due to tumors), one Fgf21 KO vehicle-treated mouse due to sudden death, one WT LA-Gcg treated mouse (euthanized due to tumors), and two LA-Gcg treated mice (one euthanized due to tumors, one died following MRI-analysis). n=4-6.

### *In vivo* glucagon or semaglutide pharmacology in diet-induced obese WT male and female mice

A third cohort of C57BL/6J male and female mice (Janvier, n=24 males, n=23 females) were maintained on a HFHS diet (Research Diets, #D12331i) from 10 weeks to 56/78 weeks of age (male/female). Before the study, single-housed males and double-housed females were grouped into blocks according to body weight (Male/female vehicle group: 56.3 g/54,0 g; semaglutide: 57 g/53 g; LA-Gcg: 57.1 g/ 53.8 g), ensuring experimental grouping with similar body weight across sex. Female and male mice (n=8 for all groups except n=7 for males injected with semaglutide) were subcutaneously injected with vehicle, 30 nmol/kg LA-Gcg (Novo Nordisk Compound Sharing, #NNC9204-0043), or 30 nmol/kg semaglutide (MedChemExpress, #HY-114118) dissolved in a vehicle solution. Food intake and body weight were monitored daily. Mice were subjected to an ipGTT on experimental day 11 as detailed below. Body composition could not be measured because the MRI equipment was being repaired at that time. The necropsy of animals was performed on experimental day 14, euthanizing animals by decapitation for collection of trunk blood.

### Intraperitoneal glucose tolerance test (ipGTT)

Mice were fasted for 4 hours in the morning and challenged with an intraperitoneal glucose load: 1.5 mg D-glucose per gram of body weight dissolved in isotonic saline at an injection volume of 5 µL/g of body weight. Blood glucose was measured from tail blood at time points 0-, 15-, 30-, 60-, and 120 minutes after the glucose load using a CONTOUR® XT glucometer.

### Analysis of body composition

Total fat and lean mass were determined using nuclear magnetic resonance (Bruker LF90II Body Composition Analyzer).

### Necropsy

All mice were euthanized by decapitation to obtain blood, and tissues were immediately snap-frozen in liquid nitrogen or collected on dry ice before storage at −70°C unless otherwise specified. For the dissection of the liver, the median lobe was used for RT-qPCR, and the left lateral lobe was immersed in formalin for histology.

### Plasma biochemistry

All plasma samples, except the ones collected after overnight fasting, were collected under non-fasted conditions in K2-EDTA-coated tubes (Sarstedt, 20.1341.100), chilled on ice, and centrifuged at 4°C for 10 minutes at 1,300 x rcf. Plasma was collected and aliquoted for storage at −70°C until use. Commercially available kits were used according to manufacturers’ protocols to quantify total plasma cholesterol (Stanbio Laboratory, #1010-430), triglycerides (Stanbio Laboratory, #2100-430), and non-esterified free fatty acids (Fujifilm Wako Chemical Europe, #434-91795, #436-91995, #270-77000). Plasma concentrations of insulin and FGF21 were determined using the sandwich ELISA principle against insulin in non-diluted plasma (Crystal Chem, #90080) and FGF21 in 1:3-diluted plasma (BioVendor R&D, #RD291108200R), respectively. A CLARIOstar® Plus Plate Reader was used to measure the absorbance in each sample using 96-well plates with a flat bottom.

### Histology

#### Preparation and imaging

Livers (left lateral lobe) were dissected and a small piece of approximately 10mm x 10mm was placed into cassettes (ActivFlo Routine I, #39LC-500-1) and submerged into buffered 4% paraformaldehyde (VWR, #9713.5000) for 24h. Samples were then paraffin-embedded, cut into sections, and stained with hematoxylin-eosin by the Core Facility for Integrated Microscopy (Department of Biomedical Sciences, University of Copenhagen). Images of slides stained with hematoxylin-eosin were obtained on a Zeiss Axio Observer microscope with an Axiocam 702 camera at 10x magnification.

#### Photoquantification of hepatic steatosis

We manually deselected portal veins, artifacts, and bile ducts and performed an automatic pixel classifier-based threshold of size, roundness, and color of pixels. A set of five thresholders was created for the experiment as there were slight differences in the stains. Thresholders were determined visually, ensuring good separation between vesicles (pixel size: 0.3426µm; color: 25, 179, 78; sigma: 6, threshold 810/820/830/840/850). All pictures were processed as Bioformat.

The area selected by the threshold was divided by the total selected area to estimate the percentage covered by macrosteatosis. The analysis was performed blinded using the image software, QuPath (Open-Source Download from https://qupath.github.io/).

### Real-time polymerase chain reaction

#### Tissue RNA-extraction

RNeasy Lipid Tissue Mini Kit (Qiagen, #74804), was used to extract RNA from tissues of ≤50 mg size. The extraction was performed according to the manufacturer’s protocol. RNA concentrations and purity were measured using a NanoDrop One/Oneᶜ Microvolume UV-Vis Spectrophotometer with 1µL of the collected total RNA solution.

#### cDNA conversion

500 ng total RNA in 9 µL RNase-free water was mixed with 4 µL of 5x FS buffer (Thermo Fisher Scientific, #18080-044), 1 µL of random primers (Sigma-Aldrich, #11034731001), and 1 µL of 0.1M DTT (Thermo Fisher Scientific, #707265ML). To anneal primers, samples were incubated at 70 °C for 3 min in a thermal cycler (Eppendorf Mastercycler Ep S Thermal Cycler). Secondly, 1 µL of dNTPs (Thermo Fisher Scientific, #R0192), 1µL of RNaseOUT™ (Invitrogen™, #10777019), 1 µL of SuperScript™ III (Invitrogen™, #12574026), and 2 µL of RNase-free water were added to each sample. After running samples for 5 min at 25°C, 60 min at 50 °C, and 15 min at 70 °C, cDNA was obtained.

#### Polymerase chain reaction

On a 384-well plate, 5 µL cDNA was loaded with 6µL of the PCR primer/enzyme mixture containing 5.5 µL Precision® PLUS qPCR Master Mix with SYBR Green (Primer design, #PPLUS-SY-20ML), 0.1 µL of 10 µM forward primer, 0.1 µL of 10 µM reverse primer, and 0.3 µL RNase-free water. RT-qPCR program was run in a LightCycler® 480 (Roche, #05015243001), with 2 minutes of incubation at 95°C followed by 45 amplification cycles at 60°C. At last, a stepwise increase in temperature from 60°C to 95°C was performed to analyze melting curves. Melting curves were quality-checked to validate primers (not shown). RNAse-free water samples were run with each primer to ensure that no contamination was present. All samples were run as doublets, averaging the Ct values for the technical replicates. The list of primers used in this study is available in Supplementary Table 2.

#### Calculations for qPCR analysis

ΔCt values were calculated relative to the housekeeping gene, *Rplp0/36b4*, using the WT vehicle-treated group to generate the ΔΔCt values, respectively of sex. Statistical testing was performed on log2FC values (−ΔΔCt) to ensure linearity and maintenance of a more normal distribution of data points compared to those of relative expression (2^-ΔΔCT).

## Results

### Metabolic phenotyping of female Fgf21 KO mice

To better understand the physiological role of FGF21 in female mice, we initially measured body weight and food intake in adult Fgf21 KO and WT chow-fed female mice (Fig. 1A). At 13 months of age (Supplementary Fig. 1A), WT mice tended to be heavier than Fgf21 KO females (Fig. 1B), reflected in differences in body composition (Fig. 1C). While total lean mass was similar between groups, Fgf21 KO mice had significantly less fat mass (Supplementary Fig. 1B). During the 37 days of phenotyping on chow diet, food intake and body weight were similar between WT and Fgf21 KO female mice (Fig. 1D, Supplementary Fig. 1C). To examine how FGF21 deficiency influences female metabolic adaptation to starvation, particularly in ketogenesis, we subjected Fgf21 KO and WT female mice to overnight fasting, a condition known to induce metabolic stress ^18^. Body weight loss and refeeding behavior after overnight fasting were similar between WT and Fgf21 KO female mice (Fig. 1E, F). Fasting induced no significant changes between genotypes in blood glucose and the ketone body beta-hydroxybutyrate (Fig. 1G, H), plasma NEFAs, cholesterol, and triglycerides (Fig. 1I-K). A second cohort of WT and Fgf21 KO female mice of similar age (Supplementary Fig. 1D) was exposed to voluntary exercise in running wheels (Fig. 1A). Voluntary running wheel activity and exercise-induced weight loss were comparable between WT and Fgf21 KO female mice (Fig. 1L, Supplementary Fig. 1E). After this voluntary running period, blood glucose and organ (kidney, liver, brown adipose tissue, inguinal white adipose tissue, and epididymal white adipose tissue) weights were similar between groups (Supplementary Fig. 1F-I). While some differences were observed in body composition at baseline, these data suggest that the overall impact of FGF21 deficiency on the physiological response to fasting and voluntary exercise in female mice exposed to a chow diet appears limited.

**Figure 1.**
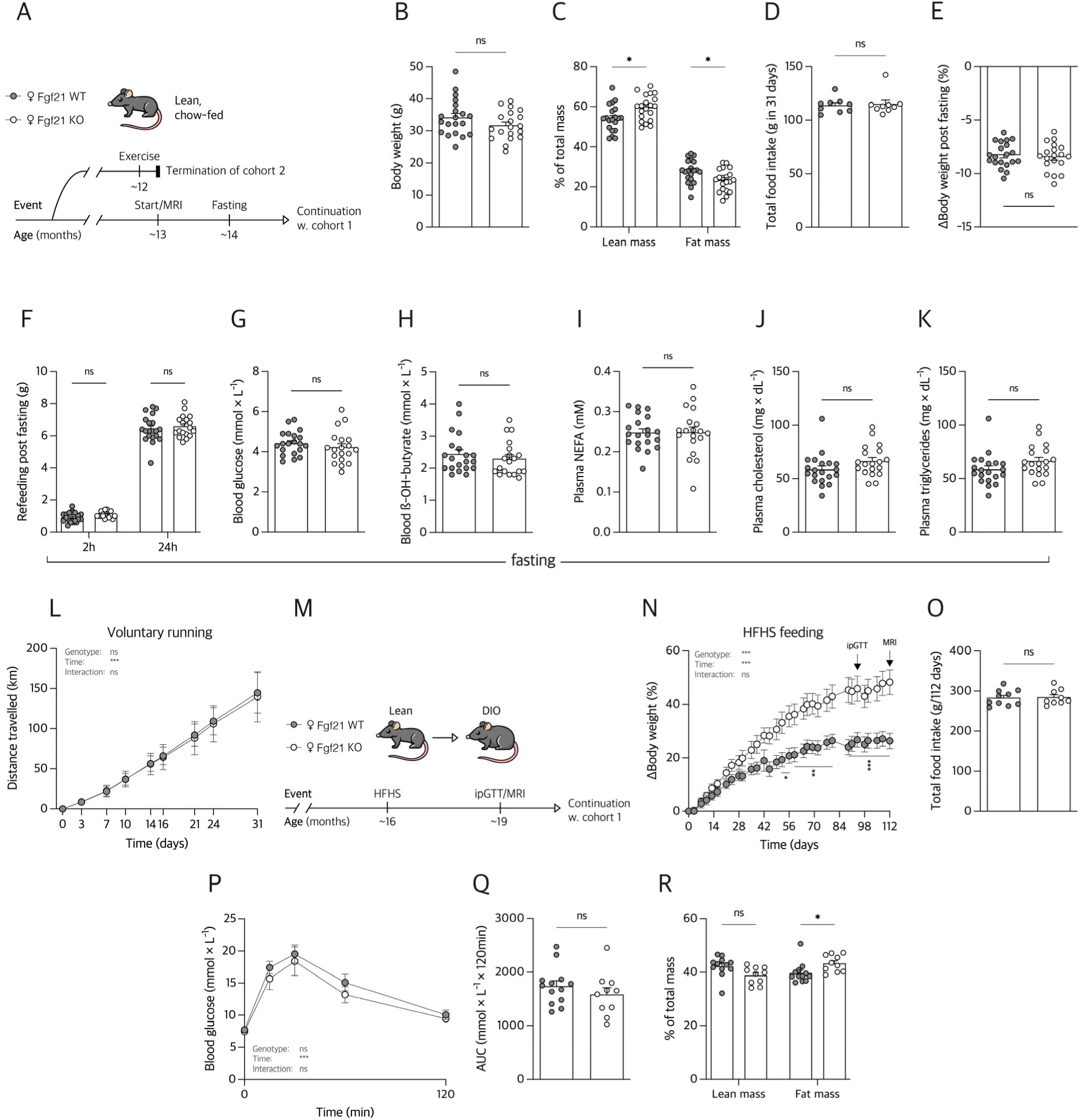
**A**, illustration of study design for panels B-L for lean chow-fed Fgf21 KO and WT female mice. **B**, total body weight, and **C**, the relative distribution of lean- and fat mass. **D**, total intake of chow diet in 31 days. **E**, body weight response to 16h of fasting, and **F**, the refeeding pattern 2h and 24h post end of fasting. **G**, blood glucose response, and **H**, blood ß-OH-byturate after 16h of fasting. Plasma measures (from tail blood) of 16h-fasted mice: **I**, non-esterified free fatty acid (NEFA); **J**, plasma cholesterol; and **K**, plasma triglycerides. **L**, cumulative distance traveled during 31 days of exposure to a running wheel. **M**, illustration of study design for panels **N-R** for the induction of diet-induced obesity (DIO) in Fgf21 KO and WT mice using a high-fat, high-sucrose diet (HFHS). **N**, the percentage change in body weight for 112 days, and **O**, the corresponding total food intake of the HFHS diet. **P**, time graph of blood glucose levels following an intraperitoneal glucose tolerance test (ipGTT) on day 94 with, **Q**, an area under the curve (AUC) assessment. **R**, distribution of lean- and fat mass following an MRI performed on day 112. n=19-20 except for B-K, n=5-7 for **L**, and n=10-13 for **N**-**R**. Data from **L**, **N**, and **P** were analyzed by two-way analysis of variance (ANOVA), with follow-up Bonferroni-corrected unpaired t-tests, to assess the effect over time. Data from **C**, **F** and were analyzed using multiple unpaired Welch t-tests, Bonferroni-corrected. Data from **B**, **D**, **E**, **G-K**, **O** and **Q** were analyzed using an unpaired Welch t-test. Bar graphs are mean ± SEM. \**P* < 0.05, \*\**P* < 0.01, \*\*\**P* < 0.001.

To investigate the role of FGF21 under obesogenic conditions in females, we exposed mice in cohort 1 to a high-fat, high-sucrose (HFHS) diet (Fig. 1M). Interestingly, we observed that Fgf21 KO female mice gained significantly more weight (+48%) than WT females (+26%) following the HFHS dietary intervention (Fig. 1N, Supplementary Fig. 1J). This enhanced weight gain was independent of changes in food intake (Fig. 1O, Supplementary Fig. 1K). Despite this increased weight gain, Fgf21 KO female mice exhibited similar glucose tolerance to WT mice (Fig. 1P, Q). After 16 weeks of HFHS feeding, Fgf21-deficient female mice showed a more significant proportion of fat mass and a trend toward lower lean mass without changes in absolute fat or lean mass (Fig. 1R, Supplementary Fig. 1L). Collectively, our data suggests, in contrast to the negligible effects of FGF21 deficiency in chow-fed female mice, that FGF21-deficient females exhibit a more obesity-prone phenotype upon exposure to an obesogenic diet.

### Pharmacological effects of a long-acting glucagon analog in diet-induced obese wild-type and Fgf21 KO female mice

To study the role of FGF21 in mediating the metabolic effects of glucagon in females, we investigated the effects of a long-acting glucagon analog (LA-Gcg) on diet-induced obese (DIO) WT and Fgf21 KO mice from cohort 1 (Fig. 2A). Daily subcutaneous injections of 30 nmol/kg LA-Gcg for 16 days resulted in a decrease in body weight in both WT and Fgf21 KO female mice (Fig. 2B, Supplementary Fig. 2A). A significant interaction effect (Treatment x Genotype, Fig. 2B, Supplementary Fig. 2A) indicated that the treatment effect on weight varied across genotypes. Neither treatment nor genotype significantly affected food intake (Fig. 2C, Supplementary Fig. 2B), suggesting that LA-Gcg-induced weight loss may be mediated by increased energy expenditure or energy excretion. While LA-Gcg impaired glucose tolerance in both WT and Fgf21 KO mice (Fig. 2D, E), the significant interaction between genotype and LA-Gcg treatment suggests that Fgf21 deficiency may exacerbate this effect in females. LA-Gcg treatment tended to reduce fat mass in WT mice, although this effect was not statistically significant (Supplementary Fig. 2C). In contrast, LA-Gcg significantly reduced lean mass in both WT and Fgf21 KO female mice (Supplementary Fig. 2D).

**Figure 2.**
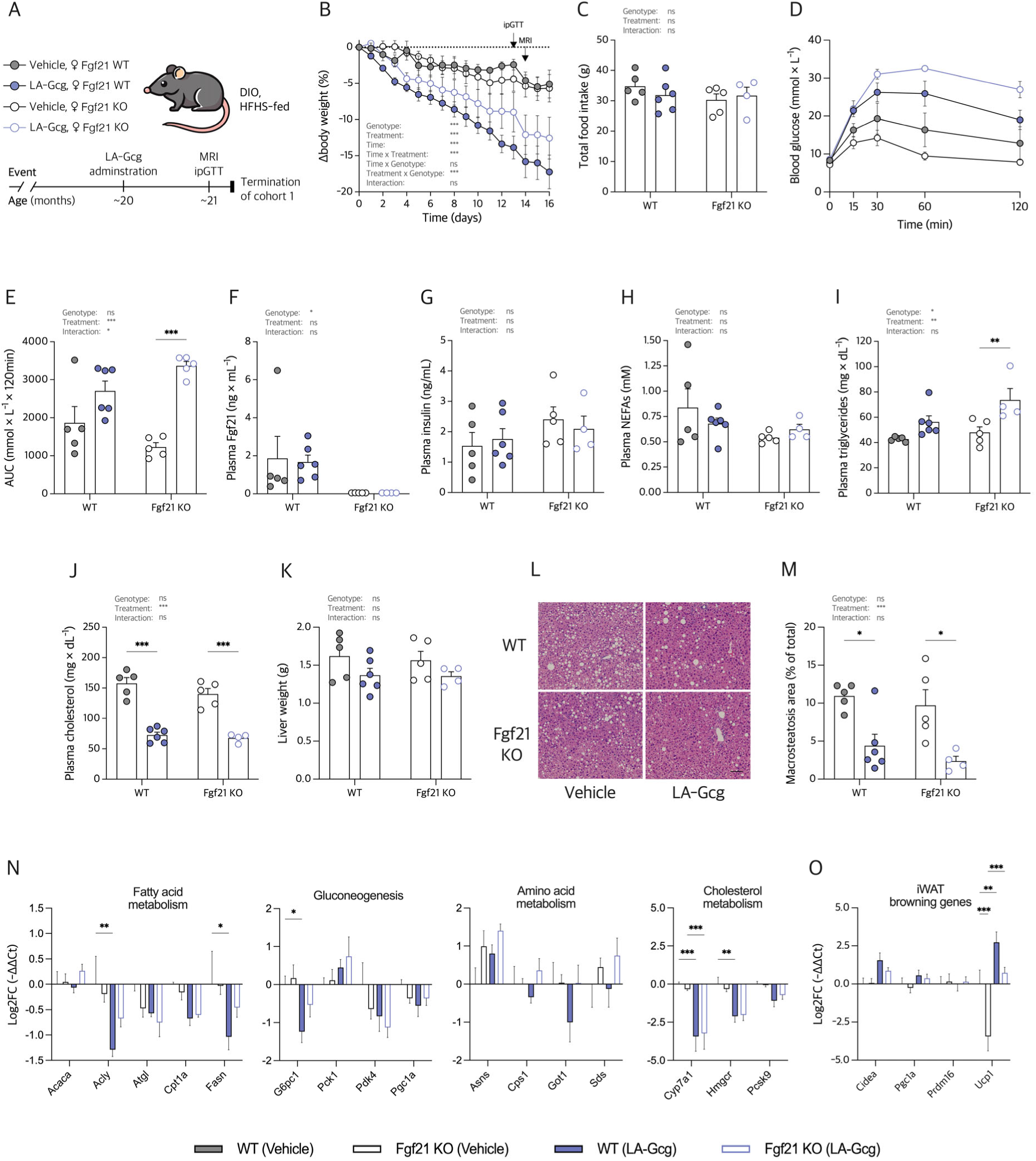
**A**, an illustration of the study design for female diet-induced obese (DIO) female mice fed with a high-fat high-sugar (HFHS) diet and treated with either vehicle or 30 nmol x kg^-1^ long-acting glucagon (LA-Gcg). **B**, time graph of % change in body weight. **C**, total food intake in 16 days. **D**, blood glucose levels following an intraperitoneal glucose tolerance test (ipGTT) on day 13 with, **E**, the corresponding area under the curve (AUC) assessment. **F**, Plasma FGF21, **G**, plasma insulin, **H**, plasma non-esterified fatty acids (NEFAs), **I**, plasma triglycerides, and **J**, plasma cholesterol levels obtained from necropsy. **K**, total liver weight. **L**, representative images of H&E-stained liver slides used for, **M**, macrosteatosis image quantification. **O**, log2 fold change (log2FC) in gene expression of liver mRNA relative to *Rplp0*/WT-Vehicle with related functions of genes displayed. **P**, log2FC in gene expression of inguinal white adipose tissue (iWAT) mRNA relative to a housekeeping gene *Rplp0* and WT-Vehicle. Data from **C**, **E-K**, and **M** were analyzed by two-way analysis of variance (ANOVA), with follow-up Bonferroni-corrected unpaired t-test, one family. A three-way ANOVA was used to analyze time graphs **B**. Multiple Bonferroni-corrected unpaired Welch t-tests were used for **N** and **O**, one family per gene. Area under the curve (AUC) from each data point in **D** was used to generate data points in **E**. Scalebar on L corresponds to 100µm. Bar- and time graphs are mean ± SEM. \**P* < 0.05, \*\**P* < 0.01, \*\*\**P* < 0.001.

While glucagon administration is known to induce FGF21 secretion in male mice^34^, we observed no increase in plasma FGF21 levels in WT female mice following LA-Gcg treatment (Fig. 2F). Neither treatment nor genotype altered plasma insulin (Fig. 2G) or NEFA levels (Fig. 2H). LA-Gcg significantly increased circulating triglycerides in Fgf21 KO mice (Fig. 2I). While genotype appeared to influence triglyceride levels, no specific group differences reached statistical significance (Fig. 2I). Finally, LA-Gcg treatment, but not genotype, decreased plasma cholesterol levels (Fig. 2J).

To determine if FGF21 deficiency altered the hepatic effects of LA-Gcg, we assessed liver weights and hepatic macrosteatosis after the intervention. LA-Gcg treatment tended to reduce liver weights (Fig. 2K), whereas the lower liver-to-body-weight ratio observed in Fgf21 KO mice may be attributable to their initially greater body weight (Supplementary Fig. 2E). LA-Gcg significantly reduced the percentage area of hepatic macrosteatosis, independent of genotype (Fig. 2L, M, Supplementary Fig. 2F), suggesting that FGF21 is not required for glucagon-mediated beneficial effects on liver steatosis. To explore whether LA-Gcg treatment differentially impacted gene expression related to key metabolic pathways in Fgf21 KO female mice, we examined the expression of genes involved in fatty acid metabolism, gluconeogenesis, amino acid metabolism, and cholesterol metabolism in the liver. In line with the reversal of hepatic steatosis, LA-Gcg treatment reduced the expression of key lipogenic genes such as *Acly* and *Fasn* (Fig. 2N). LA-Gcg treatment also targeted cholesterol metabolism, as evidenced by the downregulation of *Cyp7a1* and *Hmgcr* gene expression in WT and Fgf21 KO female mice (Fig. 2N), suggesting that LA-Gcg modulates lipogenesis and cholesterol metabolism independent of FGF21 in female mice. In inguinal white adipose tissue (iWAT), *Ucp1* gene expression was markedly reduced in vehicle-treated Fgf21 KO female mice compared to WT controls (Fig. 2O). LA-Gcg treatment restored *Ucp1* expression in Fgf21 KO expression to levels comparable to those of vehicle-treated WT mice, though not to the same extent as LA-Gcg-treated WT mice (Fig. 2O). These findings highlight the distinct effects of LA-Gcg on gene expression in liver and adipose tissue and suggest potential tissue-specific roles for FGF21 in mediating some of these effects.

### Long-acting glucagon analog versus semaglutide in diet-induced obese male and female wild-type mice

Next, we aimed to investigate the potential sex-specific effects of glucagon agonism on weight loss. To distinguish glucagon-specific actions from general weight loss mechanisms and control for sex-dependent factors, we included semaglutide, a glucagon-like peptide 1 (GLP-1) receptor agonist approved for obesity treatment, as a comparator. Female and male diet-induced obese C57BL/6J mice were treated equimolarly with 30 nmol/kg LA-Gcg or semaglutide (Fig. 3A). While semaglutide induced similar weight loss in both sexes (30% loss in females, 33% loss in males), LA-Gcg treatment resulted in significantly less weight loss in females (−16%) compared to males (−25%) (Fig. 3B, C, Supplementary Fig. 3A). LA-Gcg significantly reduced food intake in male mice, an effect not observed in females (Fig. 3D, Supplementary Fig. 3B). Semaglutide, however, similarly suppressed food intake in both sexes (Fig. 3D, Supplementary Fig. 3B). These findings demonstrate clear sex-specific effects of LA-Gcg on weight loss and food intake, distinct from the impact of semaglutide, and suggest potential sex-dimorphic responses in glucagon signaling. To gain insight into the underlying mechanisms contributing to these sex-specific differences, we also investigated parameters related to metabolic health. While semaglutide improved glucose tolerance, particularly in males (Fig. 3E, F), LA-Gcg had the opposite effect, worsening glucose tolerance in both sexes, especially in females (Fig. 3F). Despite this impaired glucose tolerance, both male and female mice treated with LA-Gcg exhibited significantly lower plasma insulin levels at termination (Fig. 3G). Semaglutide treatment also decreased plasma insulin, but this effect was significant exclusively in males (Fig. 3G). Despite a significant effect of treatment on plasma FGF21 levels and a trend toward an increase after LA-Gcg treatment, this increase was not statistically significant across groups (Fig. 3H). LA-Gcg significantly increased plasma NEFA levels, independent of sex (Fig. 3I), with a similar trend observed for plasma triglycerides (Fig. 3J). These changes may indicate increased expenditure or lipid mobilization with LA-Gcg treatment. Both treatments potently lowered plasma cholesterol compared to vehicle-treated controls (Fig. 3 K).

**Figure 3.**
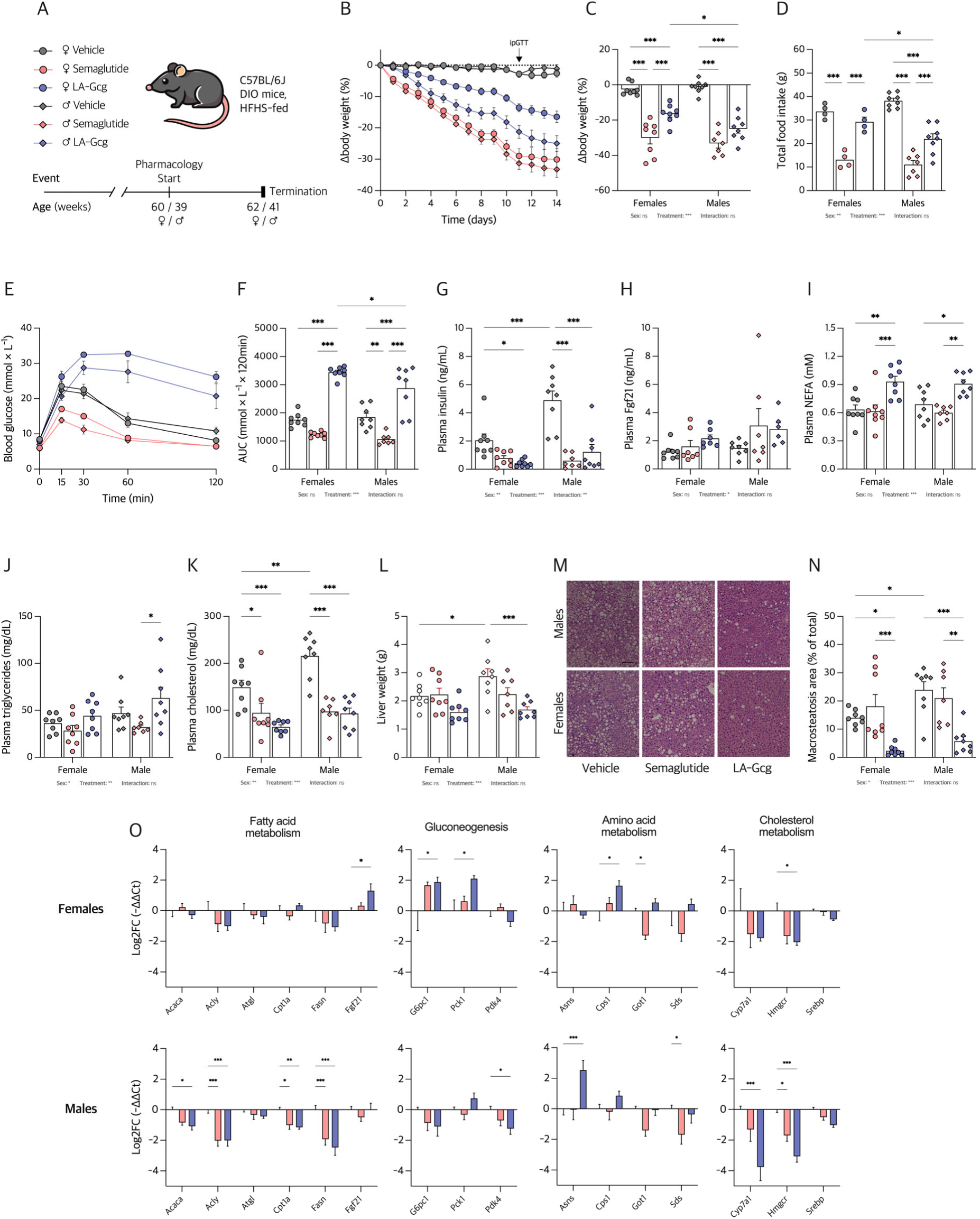
**A**, an illustration of study design using high-fat high-sugar (HFHS)-fed diet-induced obese (DIO) female and male WT mice, treated with either vehicle, 30 nmol x kg^-1^ semaglutide, or 30 nmol x kg^-1^ long-acting glucagon analog (LA-Gcg). **B**, time graph of % change in body weight resulting in, **C**, a total % change in body weight at day 14. **D**, total food intake. **E**, blood glucose levels following an intraperitoneal glucose tolerance test (ipGTT) on day 11 with, **F**, the corresponding area under the curve (AUC) assessment. **G**, plasma insulin, **H**, Plasma FGF21, **I**, plasma non-esterified fatty acids (NEFAs), **J**, plasma triglycerides, and **K**, plasma cholesterol levels obtained from necropsy. **L**, total liver weight. **M**, representative images of H&E-stained liver slides used for, **N**, macrosteatosis image quantification. **O**, log2 fold change (log2FC) in gene expression of liver mRNA relative to *Rplp0* and vehicle in female (upper panel) and male (lower panel) mice, respectively, sorted by annotated function. Data from **C**, **D**, **F-L, and N** were analyzed by two-way analysis of variance (ANOVA), with follow-up Bonferroni-corrected unpaired t-test, two families. A three-way ANOVA was used to analyze time graph **B**. ANOVA results are listed below the graphs. Multiple Bonferroni-corrected unpaired Welch t-tests were used for **O**, one family per gene. Area under the curve (AUC) from each data point in **E** was used to generate data points in **F**. Bar- and time graphs are mean ± SEM. \**P* < 0.05, \*\**P* < 0.01, \*\*\**P* < 0.001. n=8 for all groups except male semaglutide-treated mice.

Given the reported benefits of glucagon on liver health ^31,38^, we assessed the impact of the treatments on hepatic metabolic health. In LA-Gcg-treated mice, the decrease in body weight appeared to follow the decrease in liver weight (Fig. 3L). Conversely, the rapid weight loss induced by semaglutide increased the liver-to-body weight ratio in both sexes, although this increase was only significant for females (Supplementary Fig. 3C). This effect may be attributed to the depletion of intrahepatic fat observed with LA-Gcg treatment in both sexes, as evidenced by histological analysis of macrosteatosis (Fig. 3M, N, Supplementary Fig. 3D). To further elucidate the mechanisms underlying these changes in the liver, we analyzed the effects of semaglutide and LA-Gcg on the expression of genes related to fatty acid metabolism, gluconeogenesis, amino acid metabolism, and cholesterol metabolism in male and female mice (Fig. 3O). In females, LA-Gcg significantly upregulated *Fgf21*, *G6pc1*, *Pck1*, and *Cps1*, and downregulated cholesterol metabolism genes, including *Hmgcr* and *Cyp7a1*, compared to vehicle (Fig. 3O). In males, LA-Gcg had more pronounced effects across multiple pathways, significantly downregulating fatty acid metabolism genes (*Acaca*, *Acly, Cpt1a*, and *Fasn*), *Pdk4*, and cholesterol metabolism genes (*Hmgcr* and *Cyp7a1*), while upregulating *Asns* (Fig. 3O). Notably, *Fgf21* expression was largely unaffected in male mice (Fig. 3O), consistent with plasma FGF21 levels (Fig. 3H). Collectively, these findings demonstrate treatment-specific modulation of metabolic pathways, with LA-Gcg exhibiting a more potent effect, particularly in male mice, compared to semaglutide, in line with the sex-specific impact of LA-Gcg on body weight loss.

## Discussion

This study reveals a striking sexual dimorphism in the glucagon-FGF21 endocrine axis, a pathway with implications for both obesity and diabetes. While the importance of FGF21 in energy homeostasis has gained significant attention, preclinical studies, particularly those investigating its interaction with glucagon, have predominantly focused on male rodents^32–34,39^. This bias limits our understanding of FGF21 and glucagon actions in females, despite the near-equal prevalence of obesity in both sexes^8,9^ and the sexually dimorphic nature of many metabolic parameters and related comorbidities^1^. Our initial characterization of Fgf21 KO females on a chow diet revealed only subtle phenotypic differences compared to wild-type controls. However, when reproductively senescent (18-month-old) Fgf21 KO females were challenged with an obesogenic diet, they exhibited increased weight gain, highlighting a key role for FGF21 in the context of dietary stress. This observation prompted us to investigate the role of FGF21 in mediating the established beneficial effects of glucagon on body weight, a phenomenon previously described exclusively in male mice^32^. Our findings indicate that FGF21 deficiency significantly blunts the glucagon-mediated reduction in body weight and exacerbates glucose intolerance in aged female mice. Furthermore, we uncover a pronounced sex-specific response to a long-acting glucagon analog. Given the emerging use of glucagon receptor agonism and FGF21-based therapies for obesity and liver disease ^31,38,40^, this work illustrates the importance of sex as a critical factor in the study of the glucagon-FGF21 metabolic axis.

One notable finding from our study is the differential response of Fgf21 KO female mice to chow versus an obesogenic (HFHS) diet, with exacerbated weight gain observed exclusively on the HFHS diet. Several parameters remained unchanged between WT and Fgf21 KO females on a chow diet, such as fasting-induced ketogenesis and voluntary exercise, which are consistent with previous studies in males^41,42^. The exacerbated weight gain observed in HFHS-fed Fgf21 KO females may represent a novel sex-specific response, as similar studies in male mice have not reported remarkable differences in weight gain between WT and Fgf21 KO mice^43,44^. Our results leave the door open for further research to explore the impact of FGF21 deficiency in females under various dietary interventions, including ketogenic diets, and at a different life stage, such as in young females, where reproductive senescence would not confound the study results. Despite this, our study highlights the importance of FGF21 in energy homeostasis in females, particularly in modulating energy balance as a function of diet^14^. The increased weight gain in HFHS-fed Fgf21 KO female mice without corresponding changes in food intake supports a role for FGF21 in regulating energy expenditure or energy excretion^45^. Interestingly, this increased weight gain was not associated with worsened glucose homeostasis or other metabolic parameters, contrasting with previous studies linking FGF21 deficiency to impaired glucose tolerance in DIO male rodent models^46^. This discrepancy underscores the potential sex-specific roles of FGF21 in glucose homeostasis, warranting further investigation. Compensatory mechanisms in females may mitigate the metabolic consequences of FGF21 deficiency, as suggested by our data.

Our findings suggest a blunted weight loss response to subchronic glucagon receptor (Gcgr) agonism in FGF21-deficient female mice. Prior work in males has implicated FGF21, along with other factors such as FXR, in mediating some of glucagon’s beneficial metabolic effects, including weight loss and improvements in lipid metabolism^32,33^. Since glucagon still lowers body weight and improves cholesterol metabolism and hepatic steatosis in Fgf21 KO female mice, our findings suggest that other factors, such as FXR, may also play a role in mediating glucagon effects in females. It is important to acknowledge that the WT vs Fgf21 KO female mice cohort was quite old (20-21 months old) when we evaluated LA-Gcg treatment and that illness in some of these mice reduced the final sample size in some groups. This limitation in statistical power may have prevented us from drawing more definitive conclusions. Despite this, our results point to a sex-specific divergence in the glucagon-FGF21 axis, revealing a complex relationship between FGF21, glucagon signaling, and glucose homeostasis, painting a different picture in females than in males.

While we initially observed no impact of FGF21 deficiency on glucose metabolism in DIO females, LA-Gcg treatment worsened glucose tolerance in Fgf21 KO female mice, a finding not previously reported in male mice^32,47^. The glucagon-induced increased gene expression of gluconeogenic genes *G6pc1* and *Pck1* in the liver of female but not male mice (Fig. 3O) might help to understand this sexual dimorphism. In contrast with previous findings in male mice^32^, FGF21 appears dispensable for LA-Gcg-induced effects on cholesterol metabolism^15^ and hepatic lipid turnover in females. Neither cholesterol levels nor hepatic *Cyp7a1* and *Hmgcr* expression were affected by genotype. Furthermore, LA-Gcg similarly reduced hepatic steatosis across both genotypes. This effect is likely independent of weight loss, as semaglutide treatment, despite inducing comparable weight loss, did not deplete hepatic lipid droplets. Other studies in global and liver-specific FGF21-deficient male mice^32,33^ used another LA-Gcg analog concurrently with high-fat feeding in 12-week-old male mice^32^ or after 16 weeks of high-fat diet (36-week-old male mice)^33^. In contrast, we used a 16-day subchronic administration of LA-Gcg in 20-21-month-old diet-induced obese females. This difference in study design, particularly the use of aged female mice with established obesity, may explain the discrepancies in findings regarding the role of FGF21 in mediating glucagon’s effects. Interestingly, we observed *Ucp1* downregulation in iWAT of vehicle-treated Fgf21 KO females, which was partially rescued by LA-Gcg, though not to WT-treated levels. Given that both glucagon receptor and FGF21 have been implicated in increasing energy expenditure in adipose tissue^48^, these results suggest that FGF21 is also partly dispensable for glucagon-mediated *Ucp1* induction in iWAT, a proxy for thermogenesis activation. The decreased expression of *Ucp1* in iWAT of Fgf21 KO females might contribute to the differential effect of glucagon treatment in these mice.

In contrast to semaglutide, which generally improves glucose control, LA-Gcg appears to drive a distinct metabolic phenotype characterized by decreased circulating insulin and elevated glucose levels, despite both treatments promoting weight loss in female mice. These results replicate the observation found by chronic LA-Gcg administration in DIO male mice^32^, a finding that contrasts with the widely held view of glucagon as a potent stimulator of insulin secretion. The exacerbation of glucose intolerance with LA-Gcg treatment aligns with glucagon’s known role in stimulating hepatic glucose output^15^. However, our data reveal a more pronounced effect in females, suggesting potential sex-specific differences in hepatic sensitivity to glucagon receptor agonism or downstream signaling pathways. This observation echoes findings from a recent study in humans with type 2 diabetes, which reported sex differences in plasma glucagon levels during mixed-meal and glucose tolerance tests^49^, despite the different entry points (LA-Gcg administration versus endogenous glucagon secretion). The mechanisms underlying this phenotype with subchronic LA-Gcg administration warrant further investigation, particularly given the growing interest in dual and tri-agonist therapies for obesity management that leverage Gcgr agonism^31^.

In conclusion, our study reveals that FGF21 deficiency exacerbates diet-induced obesity in reproductively senescent female mice, while also demonstrating sex-dependent effects of pharmacological glucagon agonism. Specifically, LA-Gcg treatment in Fgf21 KO females resulted in blunted weight loss and exacerbated glucose intolerance compared to wild-type females, but similar beneficial effects on cholesterol metabolism and hepatic steatosis, suggesting that FGF21 may mediate some, but not all, of glucagon’s beneficial effects in females. Furthermore, LA-Gcg treatment elicited a blunted weight loss response and worsened glucose tolerance in females compared to males overall. These findings underscore the critical importance of considering sex as a biological variable in glucagon and FGF21 research, particularly when developing glucagon-based therapies.

## Acknowledgements

## Acknowledgments

NNC9204-0043 was provided by Novo Nordisk Compound Sharing

## Author Contributions

**Christoffer Merrild:** Formal analysis, Conceptualization, Investigation, Visualization, Writing – original draft, Writing – review & editing. **Valdemar B. I. Johansen:** Investigation, Conceptualization, Writing – review & editing. **Christoffer Clemmensen:** Conceptualization, Funding acquisition, Project administration, Supervision, Writing – review & editing. **Pablo Ranea-Robles:** Data curation, Investigation, Methodology, Conceptualization, Supervision, Visualization, Writing – original draft, Writing – review & editing.

## Guarantor Statement

C.C. and P.R-R. are the guarantors of this work and, as such, have full access to all the data in the study and take responsibility for the integrity of the data and the accuracy of the data analysis.

## Conflict of Interest Statement, Funding, and Prior Presentation

C.C. is a co-founder of Ousia Pharma. C.M. is supported by a PhD grant from the Danish Diabetes and Endocrine Academy, which is funded by the Novo Nordisk Foundation (grant number NNF22SA0079901). P.R.R was supported by a postdoctoral fellowship from The Novo Nordisk Foundation Center for Basic Metabolic Research (International Postdoctoral Fellowship). The Novo Nordisk Foundation Center for Basic Metabolic Research is an independent research center, based at the University of Copenhagen, Denmark, and partially funded by an unconditional donation from the Novo Nordisk Foundation (www.cbmr.ku.dk) (grant numbers NNF18CC0034900 and NNF23SA0084103).

## Tables

**Table 1:**
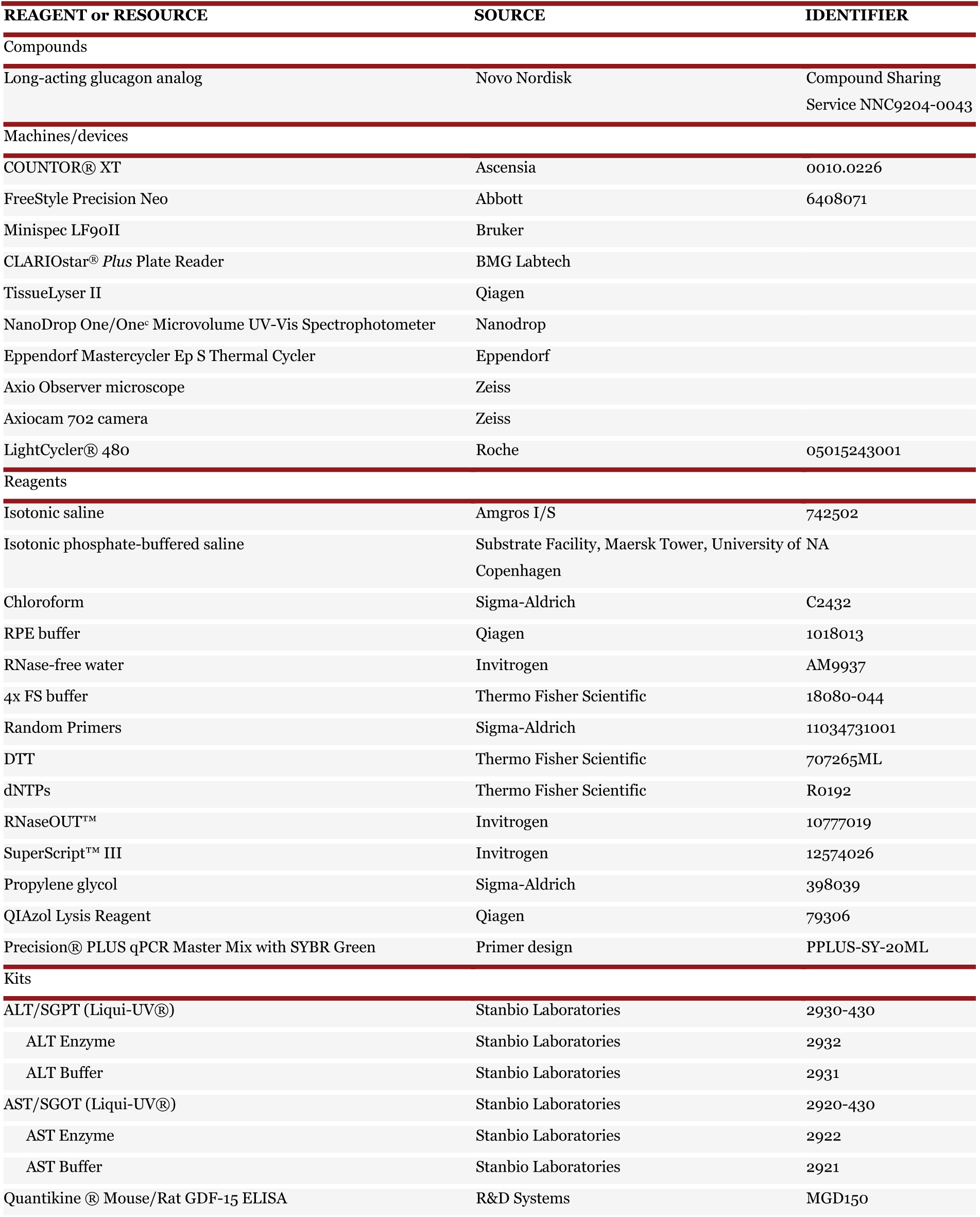
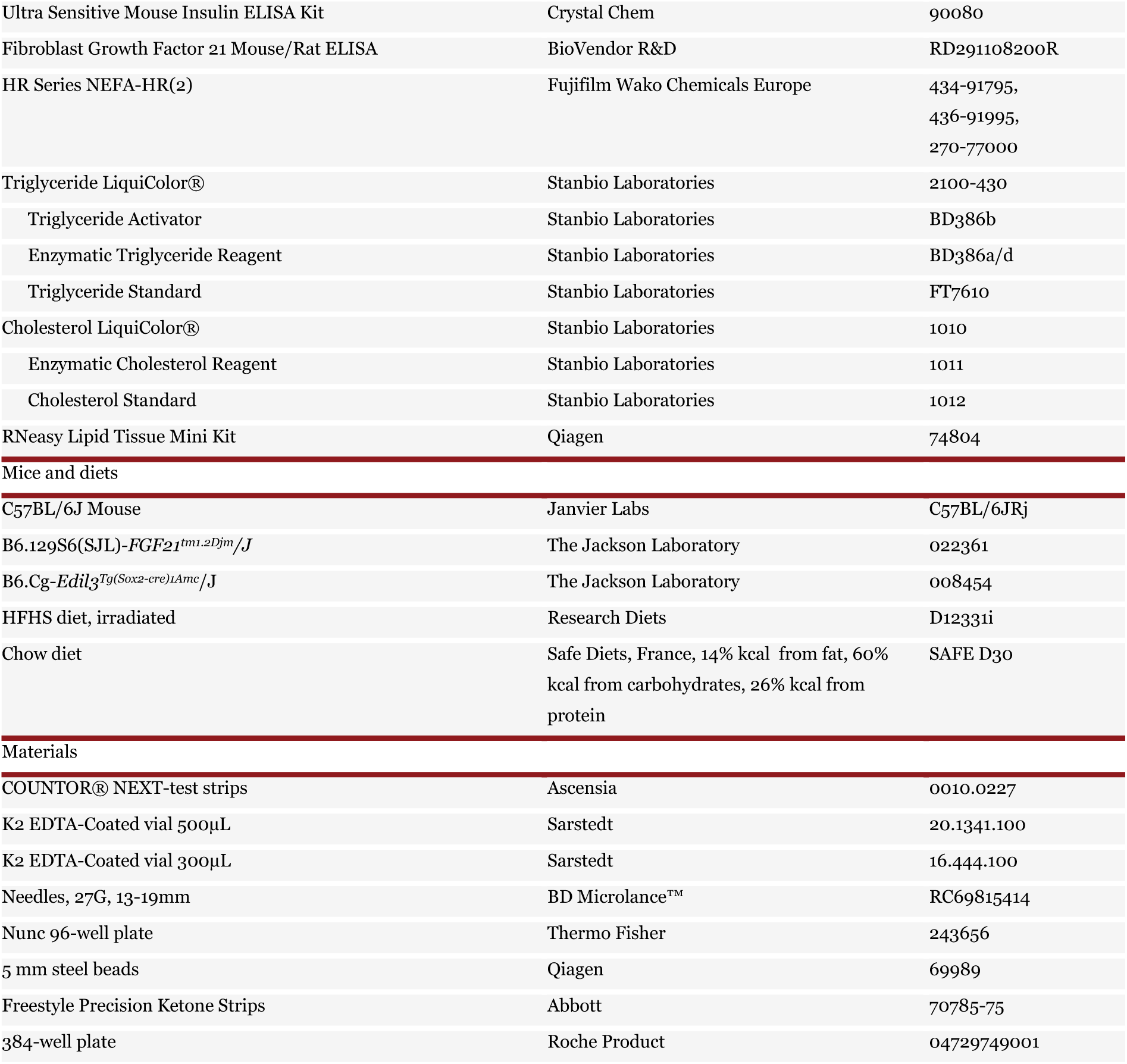
Materials.

**Table 2:**
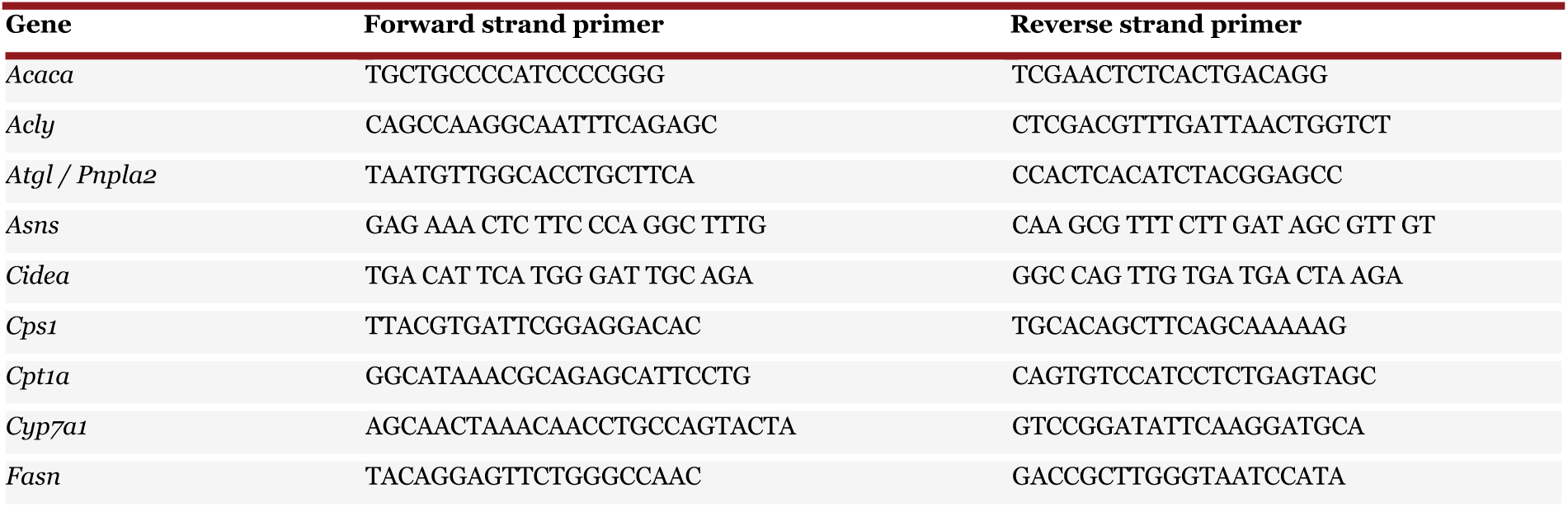
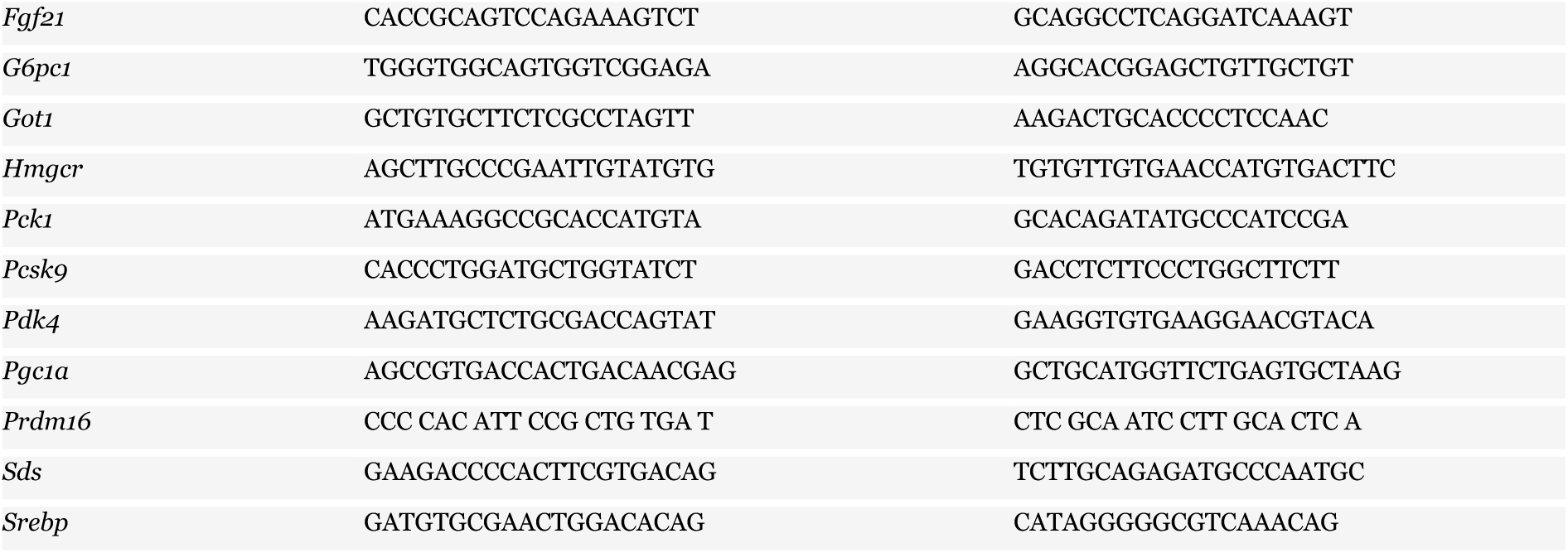
Primer sequences for q-PCR.

## Figure Legends

**Supplementary Figure 1.**
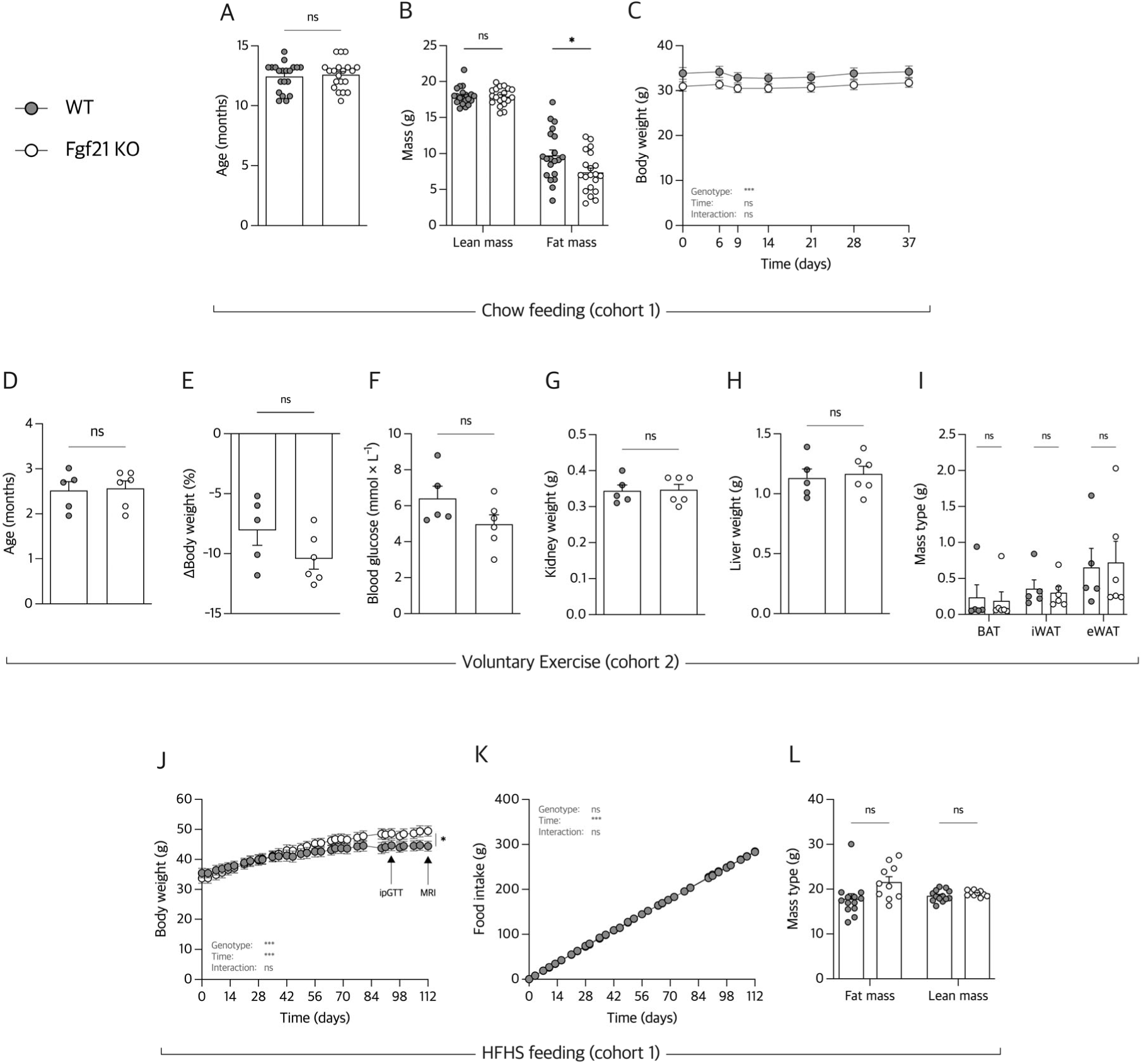
Symbols and their according groups are illustrated in the top left corner for all panels using female Fgf21 KO and WT mice. **A**, age of mice at the start of phenotyping. **B**, the total distribution of lean- and fat mass. **C**, time-resolved absolute weight changes in 37 days. **D**, age of mice at the beginning of running wheel experiment. **E**, body weight change in percentage following voluntary running for 37 days, and the corresponding, **F**, blood glucose levels. Weight of, **G**, kidney; **H**, liver; and **I**, brown adipose tissue (BAT), inguinal white adipose tissue (iWAT), and epididymal white adipose tissue (eWAT). **J**, absolute body weight as a function of time with corresponding, **K**, cumulative food intake in grams. **L**, distribution of total fat- and lean mass from MRI on day 112. n=19-20 for **A-C**, n=6-7 for **D-I**, and n=10-13 for **J-L**. Data from **C**, **J**, and **K** were analyzed by two-way analysis of variance (ANOVA), with follow-up Bonferroni-corrected unpaired t-tests, to assess the effect over time. Data from **A+D-H** were analyzed using unpaired Welch t-tests. Data from **B**, **I**, and **L** were analyzed using multiple unpaired Welch t-test, Bonferroni-corrected. Bar graphs are mean ± SEM. \**P* < 0.05, \*\**P* < 0.01, \*\*\**P* < 0.001.

**Supplementary Figure 2.**
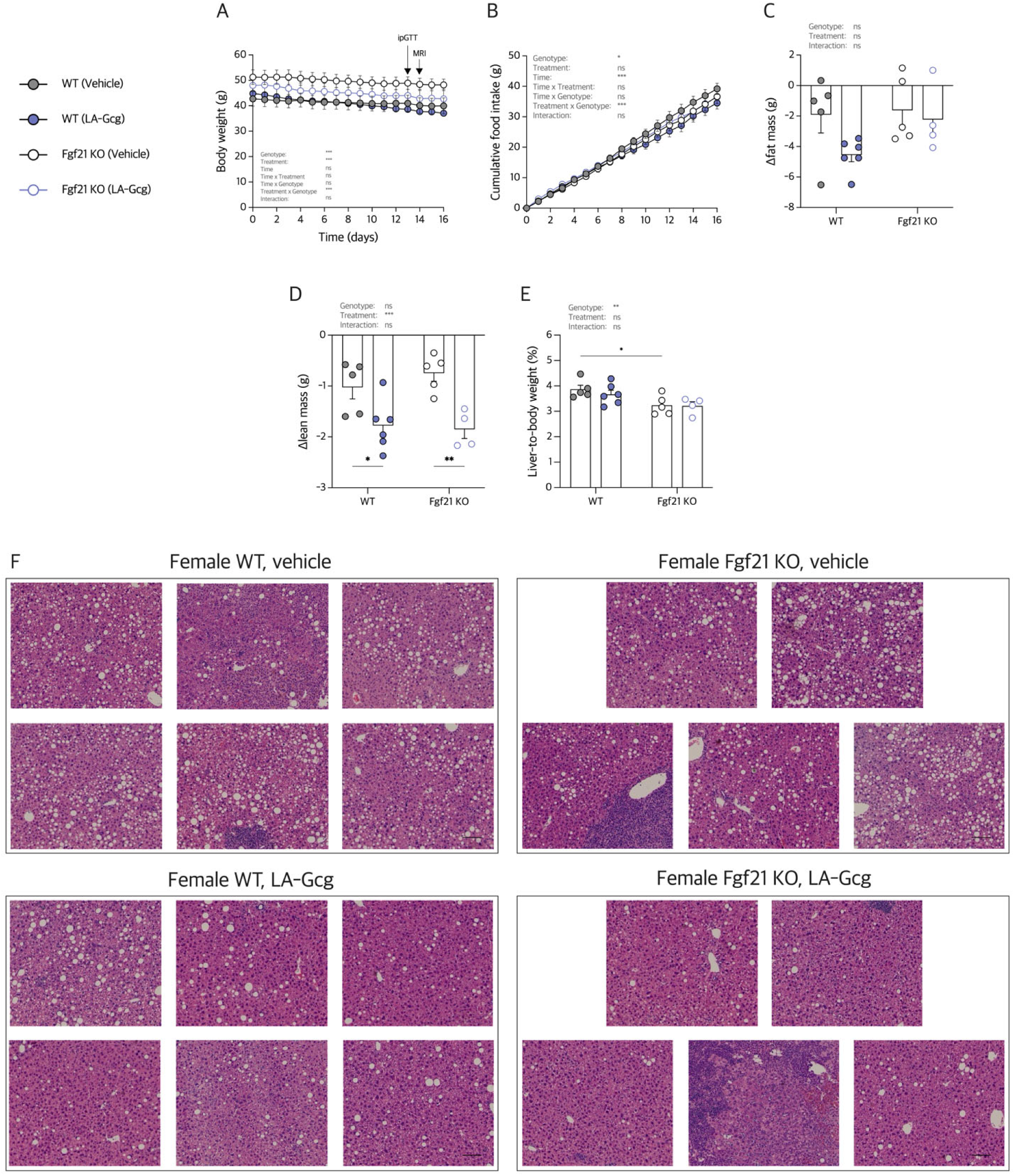
Symbols and their according groups are illustrated in the top left corner for all panels using female diet-induced obese (DIO) female mice fed with a high-fat high-sugar (HFHS) diet and treated with either vehicle or 30 nmol x kg^-1^ long-acting glucagon (LA-Gcg). **A**, time graph of total body weight. **B**, cumulative food intake in grams. **C**, total change in fat mass, and, **D**, total change in lean mass from MRI prior to study start and day 14. **E**, liver-to-body weight ratio at necropsy. **F**, images of H&E-stained liver histology. Data from **C-E** were analyzed by two-way analysis of variance (ANOVA), with follow-up Bonferroni-corrected unpaired t-test, one family. A three-way ANOVA was used to analyze time graphs **A** and **B**. Scalebar on **F** corresponds to 100µm. Bar- and time graphs are mean ± SEM. \**P* < 0.05, \*\**P* < 0.01, \*\*\**P* < 0.001.

**Supplementary Figure 3.**
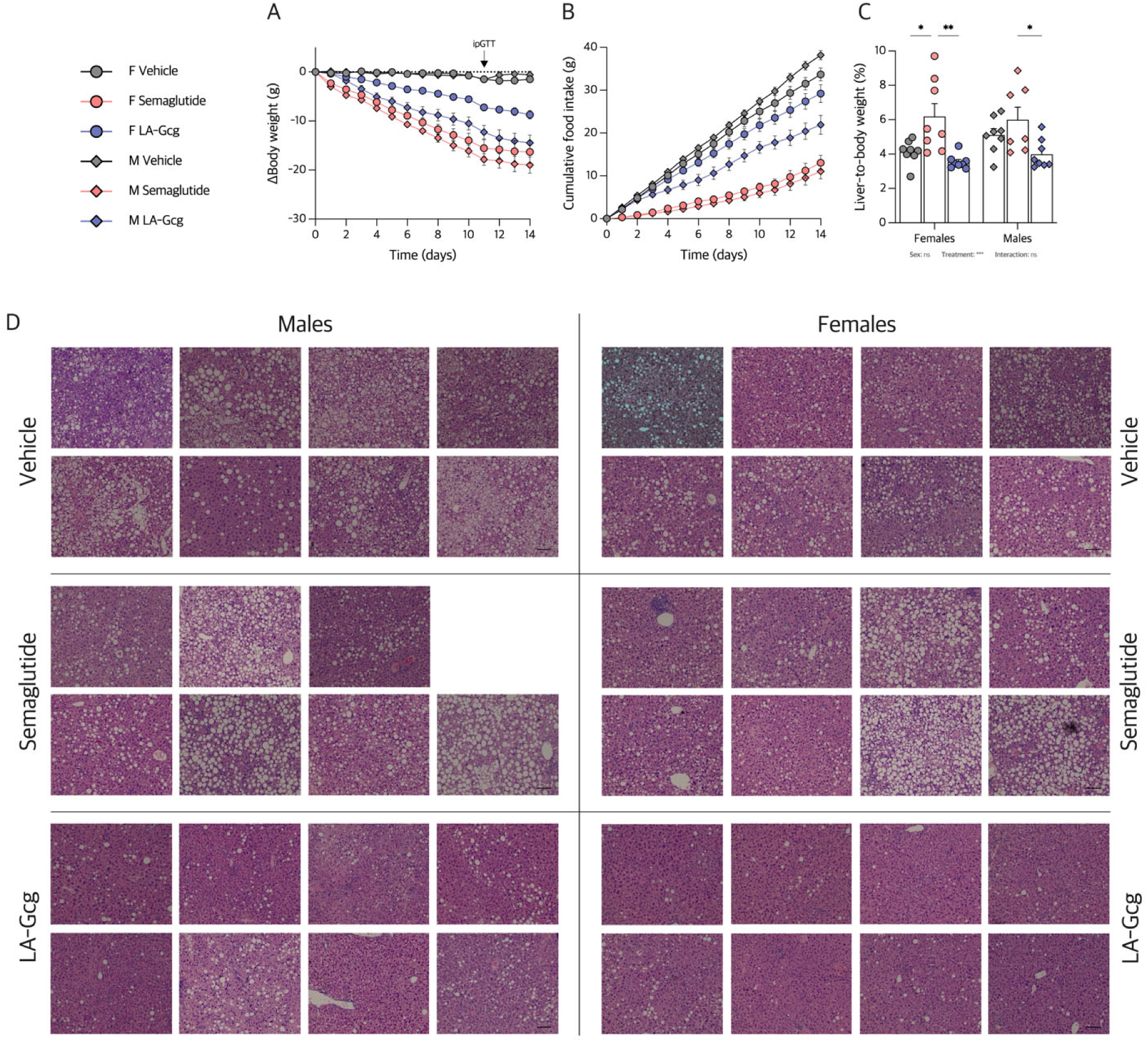
Symbols and their according groups are illustrated in the top left corner for all panels using high-fat high-sugar (HFHS)-fed diet-induced obese (DIO) female- and male WT-mice, treated with either vehicle, 30 nmol x kg^-1^ semaglutide, or 30 nmol x kg^-1^ long-acting glucagon analog (LA-Gcg). **A**, time graph of total body weight. **B**, cumulative food intake. **C**, liver-to-body weight ratio at necropsy. **D**, images of H&E-stained liver histology. Data from **D** was analyzed by two-way analysis of variance (ANOVA), with follow-up Bonferroni-corrected unpaired t-test, one family. Scalebar on E corresponds to 100µm. Bar- and time graphs are mean ± SEM. \**P* < 0.05, \*\**P* < 0.01, \*\*\**P* < 0.001. n=8 for all groups except male semaglutide-treated mice.

## Abbreviations

ANOVA: Analysis of variance
AUC: Area under the curve
BHB: Beta-hydroxybutyrate
DIO: Diet-induced obese
FGF21: Fibroblast Growth Factor 21
Gcg: Glucagon
Gcgr: Glucagon receptor
HFHS: High-fat high-sugar
ipGTT: Intraperitoneal glucose tolerance test
iWAT: Inguinal white adipose tissue
KO: Knockout
LA-Gcg: Long-acting glucagon
Log2FC: Log2 fold change
NEFA: Non-esterified fatty acid
SEM: Standard error of the mean
WT: Wildtype

